# Structural basis of GABA transport by the vesicular inhibitory amino acid transporter VIAAT

**DOI:** 10.64898/2026.09.24.754290

**Authors:** Jiahui Chen, Nanhao Chen, Jingpeng Ge, Jie Yu

## Abstract

γ-Aminobutyric acid (GABA) and glycine mediate fast synaptic inhibition by restraining neuronal excitability and stabilizing neural circuits. The vesicular inhibitory amino acid transporter (VIAAT/VGAT) loads both GABA and glycine into synaptic vesicles, but its substrate-recognition and transport mechanisms remain unclear. Here we report cryo-EM structures of human VIAAT in apo luminal-open and GABA-bound occluded states. VIAAT adopts an 11 transmembrane helix LeuT-like fold, distinct from other vesicular neurotransmitter transporters. In the central pocket, an aromatic cage comprising Phe133, Tyr221, Phe315 and Phe322 mediates recognition of GABA. Structural comparison reveals that Phe315 acts as a luminal gate, rotating away from the central pocket to open the luminal pathway for substrate release. Complemented by functional neuronal rescue experiments and a cytosolic-facing model, our findings elucidate the mechanism for VIAAT-mediated GABA transport and provide a framework for understanding how epilepsy-associated VIAAT variants impair inhibitory neurotransmission.

**Significance:** Inhibitory neurotransmission is essential for balancing brain activity, and its dysfunction leads to epilepsy and other neurological disorders. VIAAT is the sole transporter responsible for packing the inhibitory neurotransmitters GABA and glycine into synaptic vesicles, yet how it recognizes and transports these molecules has remained unknown. Here, we determine the atomic structures of VIAAT in two key states, revealing how it captures GABA and how a critical gate controls its release. These findings provide a structural framework for understanding inhibitory neurotransmission and offer a basis for deciphering how genetic mutations linked to epilepsy impair VIAAT function.

## Introduction

Fast synaptic inhibition in the vertebrate nervous system is mediated primarily by γ-aminobutyric acid (GABA) and glycine, which constrain neuronal excitability and maintain circuit stability(1, 2). Sustained inhibitory neurotransmission requires rapid refilling of synaptic vesicles with these neurotransmitters following endocytosis(3). The vesicular inhibitory amino acid transporter (VIAAT), also known as the vesicular GABA transporter (VGAT), is the only vesicular transporter identified to date that loads both GABA and glycine into synaptic vesicles (4–8). Genetic ablation of VIAAT in mice markedly reduces vesicular release of GABA and glycine at inhibitory synapses and results in embryonic lethality(7, 9). Moreover, missense variants in VIAAT have been linked to epilepsy and neurodevelopmental impairment, underscoring the essential role of VIAAT in inhibitory neurotransmission(10, 11).

VIAAT, encoded by *SLC32A1*, is the sole identified member of this transporter family and functions as a proton-coupled antiporter(5, 12). It belongs to the amino acid-polyamine-organocation (APC) superfamily and adopts a canonical LeuT-like fold(13), distinguishing it from other vesicular neurotransmitter transporters including vesicular glutamate transporters (VGLUT, SLC17)(14) and vesicular monoamine/acetylcholine transporters (VMAT/VAChT, SLC18)(15–17), which instead exhibit major facilitator superfamily (MFS) architectures. This structural divergence suggests that VIAAT may employ a transport mechanism distinct from that of other vesicular neurotransmitter transporters(18). Indeed, VGLUT transports the negatively charged neurotransmitter glutamate primarily by exploiting the vesicular membrane potential, whereas VMAT and VAChT predominantly rely on the vesicular pH gradient to drive uptake of positively charged monoamines and acetylcholine(19, 20). In contrast, VIAAT is thought to utilize both the membrane potential and the pH gradient to transport the zwitterionic neurotransmitters GABA and glycine(21), further underscoring its unique transport mechanism. However, the molecular basis of its substrate recognition and transport remains poorly understood due to the lack of atomic structural information.

Given its fundamental role in inhibitory neurotransmission, VIAAT offers limited opportunities for direct therapeutic intervention. Nevertheless, pharmacological modulators of VIAAT could serve as valuable probes for visualizing inhibitory nerve terminals in vivo in the human brain and for tracking disease progression in neurodegenerative disorders marked by GABAergic neuron loss, such as Parkinson’s and Huntington’s diseases(5, 22). To date, no VIAAT-directed modulators with high selectivity and potency have been developed. Elucidating the structural basis of substrate recognition and transport by VIAAT may help advance the development of such tools.

In this study, we determined the cryo-electron microscopy (cryo-EM) structures of VIAAT in the apo state and in complex with substrate GABA. Both structures adopt luminal-facing conformations, while substrate binding induces local conformational rearrangements that occlude the luminal translocation pathway. Supported by evidence from electrophysiological recordings in mouse primary hippocampal neurons, we identified critical residues involved in substrate recognition and transport gating. Furthermore, guided by a cytosolic-facing model, we propose a mechanism for VIAAT-mediated GABA transport into synaptic vesicles. This work establishes a structural framework for understanding proton-coupled inhibitory neurotransmitter loading, and helps explain how epilepsy-associated VIAAT variants disrupt transporter function.

## Results

### Function and structure determination of VIAAT

Wild-type (WT) full-length human VIAAT with a C-terminal Strep tag, was expressed in HEK293S GnTI^−^ cells and purified in the presence of LMNG for structural studies. Size-exclusion chromatography showed a sharp, monodisperse peak consistent with a monomeric state (SI Appendix, Fig. S1A). Notably, SDS-PAGE revealed multiple bands (SI Appendix, Fig. S1B), likely reflecting partial N-terminal degradation, consistent with the high flexibility and predicted disorder of the VIAAT N-terminal region(23). To assess the role of VIAAT in inhibitory neurotransmission, we knocked down endogenous VIAAT in cultured hippocampal neurons using short hairpin RNA (shRNA) (SI Appendix, Fig. S2), which abolished miniature inhibitory postsynaptic currents (mIPSCs) evoked by vesicular GABA release (Fig. 1A). Re-expression of human VIAAT via lentiviral transduction fully restored these currents, enabling functional analysis of VIAAT variants in the rescue neurons.

**Fig. 1.**
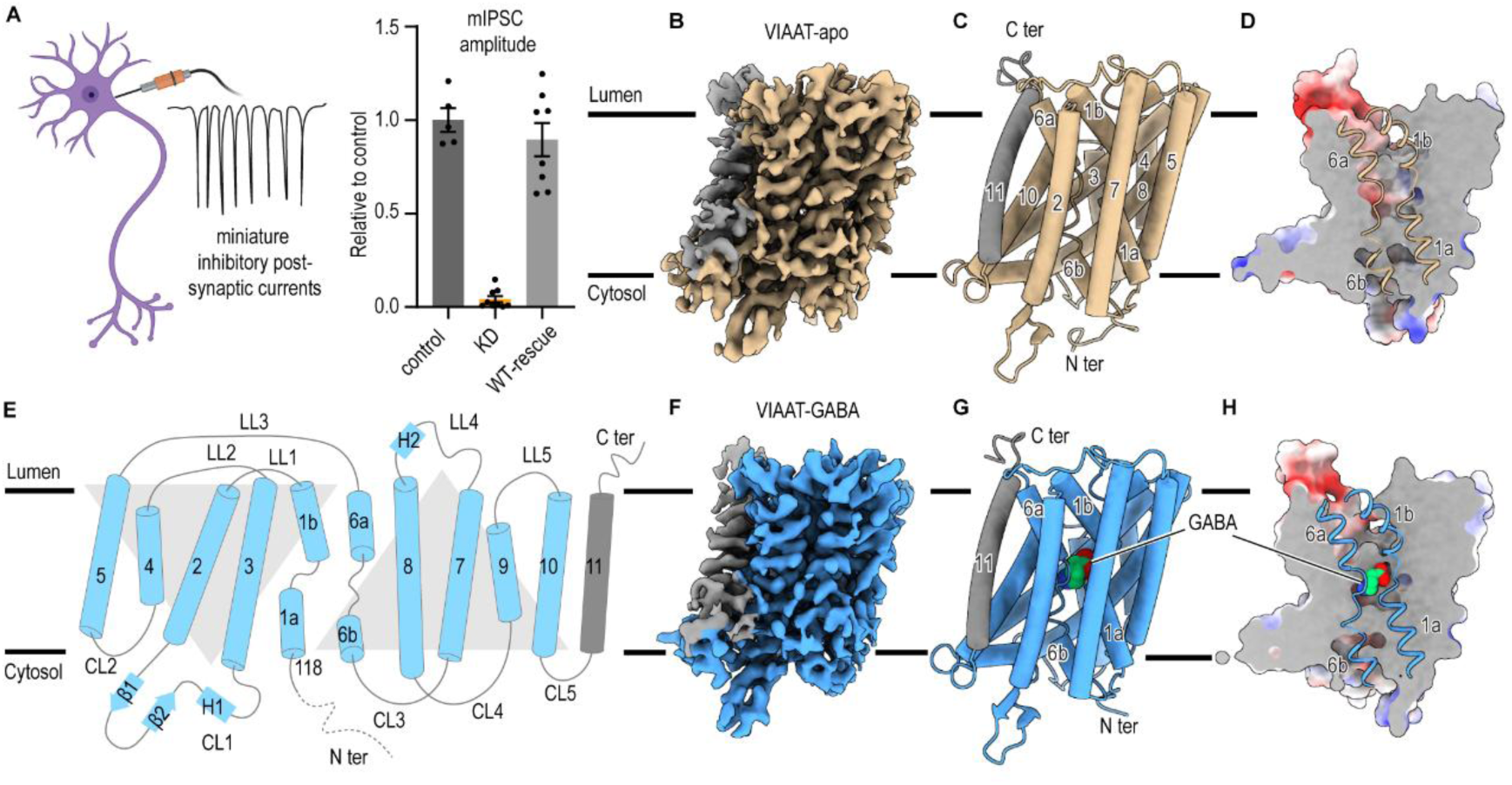
Function and structure determination of human VIAAT. **(A)** Left, schematic illustration of mIPSC recording from mouse hippocampal neurons. Right, synaptic transmission is impaired by knockdown (KD) of endogenous VIAAT and rescued by expression of wild-type (WT) human VIAAT. mIPSC amplitudes were normalized to those of the control condition (n = 5-9 cells per condition). Data are presented as mean ± s.e.m. **(B-C)** Cryo-EM map (**B**) and coordinate (**C**) of VIAAT in the apo form. TM11 is highlighted with grey. The contour level is 0.35. **(D)** Slice view of VIAAT-apo. TM1 and TM6 are shown as cartoon representations. VIAAT-apo adopts a luminal-open conformation. **(E)** Schematic representation of the structural arrangement of VIAAT. The first ten transmembrane helices (TMs) are folded into a characteristic twofold LeuT-like pseudosymmetry, consisting of an inverted-topology repeat of five TMs. TM1-TM10 are colored cyan, and TM11 is colored grey. **(F-G)** Cryo-EM map (**F**) and coordinate (**G**) of VIAAT in complex with GABA (VIAAT-GABA). TM11 is highlighted with grey. The contour level is 0.35. **(H)** Slice view of VIAAT-GABA showing the GABA-binding pocket in the central cavity. TM1 and TM6 are shown as cartoon representations. VIAAT-GABA adopts a luminal-occluded conformation.

We determined the structures of VIAAT in the apo state (VIAAT-apo) and in complex with its substrate GABA (VIAAT-GABA) at resolutions of 3.25 Å and 3.3 Å, respectively (Fig. 1B-C and F-G, SI Appendix Table 1). The large cytosolic N-terminal region was not resolved due to its highly intrinsic flexibility. Well-defined side-chain densities across most of the protein enabled accurate atomic model building (SI Appendix, Fig. S3 and S4). VIAAT comprises 11 transmembrane helices (TMs), with the N terminus facing the cytosol and the C terminus oriented toward the lumen(24) (Fig. 1C, E and G). In contrast to other vesicular neurotransmitter transporters, which adopt MFS folds, VIAAT adopts a canonical LeuT-like fold formed by two inverted pseudo-symmetric transmembrane repeats (TM1-TM5 and TM6-TM10), suggesting that VIAAT may operate through a distinct proton-coupled transport mechanism. The C-terminal TM11 of VIAAT flanks the periphery of TM2 and TM10. TM1 and TM6, which line the substrate translocation pathway, are interrupted by unwound regions that divide each helix into two segments (TM1a/1b and TM6a/6b) and constitute key structural elements for substrate recognition and binding (Fig. 1C, E). Although LeuT-like folds have been observed in many transporters, a Dali search identified drSLC38A9 from *Danio rerio* as the closest structural homolog of VIAAT among currently resolved structures(25). In both VIAAT-apo and VIAAT-GABA, the cytosolic permeation pathway is closed, indicating that these structures represent luminal-facing conformations (Fig. 1D, H).

### VIAAT in the apo state

In the VIAAT-apo structure, the cytosolic gate is occluded by tight packing among TM1a, TM3, TM6, TM8, and cytosolic loop 1 (CL1), which connects TM2 and TM3, such that the permeation pathway is accessible only from the luminal side. Particularly, Phe322 in TM6 forms hydrophobic interactions with Leu214, Leu396, and Ala400 that block solvent access to the substrate binding site at the center of transporter (Fig. 2B). To assess the functional role of Phe322, we measured mIPSCs in neurons expressing the Phe322Ala mutant in the VIAAT rescue system. The mIPSC amplitudes were markedly reduced, indicating that Phe322 is critical for transporter gating (Fig. 2D). Consistent with our findings, the Phe322Cys mutation has been reported to cause epilepsy with impaired neurodevelopment(11).

**Fig. 2.**
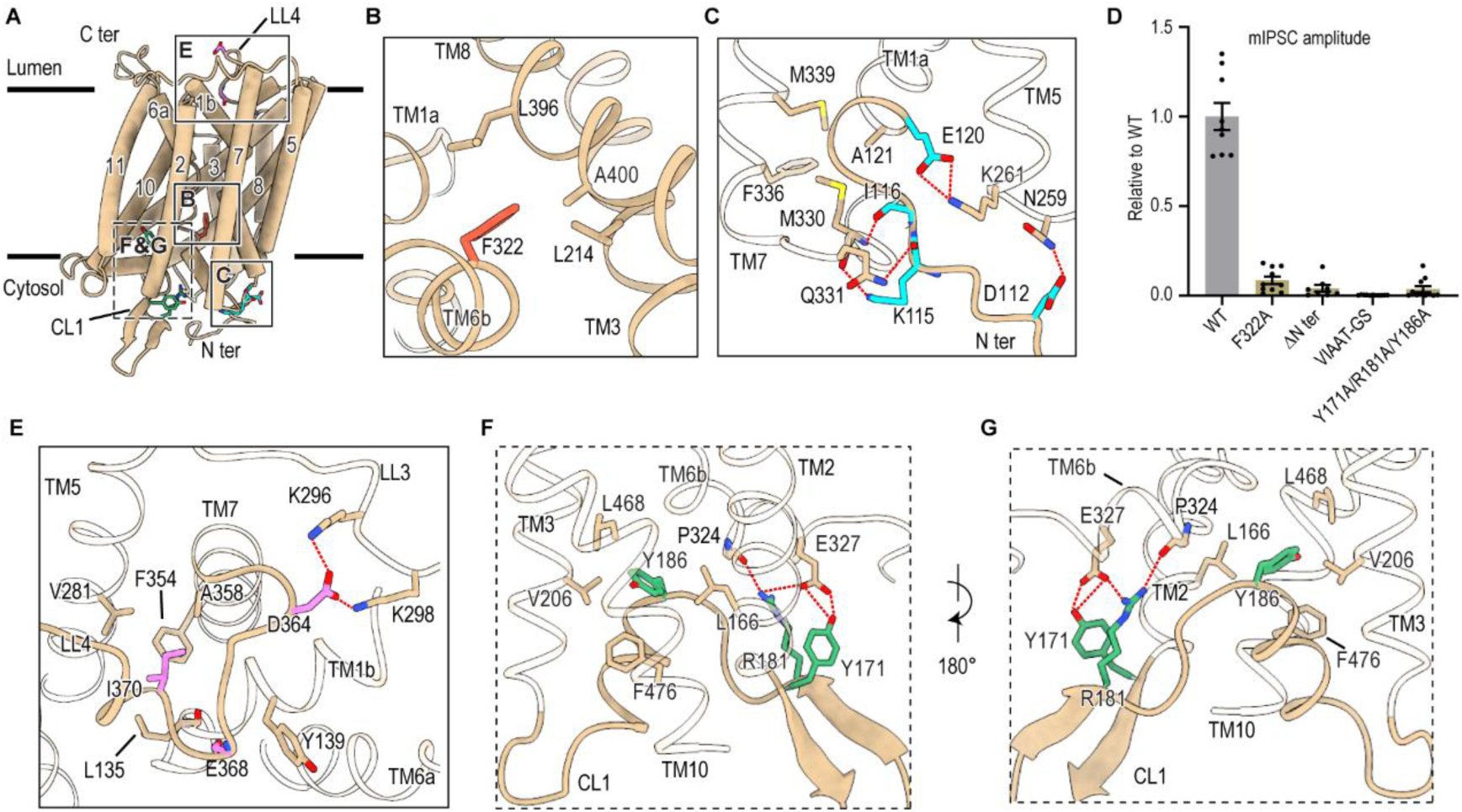
The N terminal, luminal and cytosolic loops contribute to VIAAT function. **(A)** Overall structure of VIAAT shown as a cartoon. The solid and dashed boxes in correspond to the insets in **B-C** and **E-G**. **(B)** Enlarged view of the boxed region in (**A**), showing the hydrophobic interaction network in the cytosolic side. **(C)** Enlarged view of the boxed region in (**A**), showing the interaction network at the N-terminal loop. Key residues (Asp112, Lys115, Ile116, and Glu120) in the N-terminal loop are highlighted in cyan. Hydrogen bonds are indicated by red dashed lines. **(D)** mIPSCs recorded from hippocampal neurons after knockdown of endogenous VIAAT and rescue with WT human VIAAT or its mutants. mIPSC amplitudes were normalized to those of VIAAT-WT (n = 8-11 cells per condition). Data are shown as mean ± s.e.m. **(E)** Enlarged view of the boxed region in (**A**), showing luminal loop 4 (LL4) inserting into the translocation pathway. Key residues (Asp364, Glu368, and Ile370) in LL4 are highlighted in violet. Hydrogen bonds are indicated by red dashed lines. **(F-G)** Enlarged views of the boxed region in (**A**), showing cytosolic loop 1 (CL1) wedging between TM2, TM3, TM6b, and TM10 in two different orientations. Key residues (Tyr171, Arg181, and Tyr186) in CL1 are highlighted in green. Hydrogen bonds are indicated by red dashed lines.

While most of the N-terminal region was not resolved in the VIAAT-apo structure, discernible density at the N terminus enabled partial modeling of an N-terminal loop. Residues Asp112, Lys115, Ile116, and Glu120 engage in multiple interactions with TM5 and TM6. Specifically, Asp112 forms a potential salt bridge with Asn259, and Lys115 forms two possible salt bridges with the side chain of Gln331 and the backbone carbonyl of Met330. In addition, the main chain of Ile116 forms a potential hydrogen bond with Asn331, and the side chain of Glu120 engages in possible salt bridges with Lys261 (Fig. 2C). Notably, the Met330Thr mutation has been identified in patients with genetic epilepsy(10). Given that Met330 participates in hydrophobic interactions with Phe336, Met339, and Ala121, we speculate that its hydrophobic side chain contributes to transporter stability (Fig. 2C). Although the partially resolved N-terminal region does not obstruct the cytosolic permeation pathway in our structures, previous studies of other LeuT-like transporters have shown that the N terminus can occlude the cytosolic gate and play an important role in transport(26–28). Particularly, in drSLC38A9, the N-terminal region folds into a β-hairpin that inserts into the cytosolic vestibule and exerts an inhibitory effect on transport(29). To probe the functional contribution of the N terminus, we deleted residues 1-117 (ΔN ter) and measured mIPSCs. This truncation resulted in a dramatic reduction in mIPSC amplitude (Fig. 2D), indicating that the N terminus is essential for VIAAT function. We therefore propose that the largely disordered N-terminal region may act as a dynamic cap that modulates solvent access to the central cavity of the transporter. Further studies will be required to elucidate the precise role of the N terminus in VIAAT transport.

### Luminal and cytosolic loops

Upon closer structural inspection, luminal loop 4 (LL4), which connects TM7 and TM8, and cytosolic loop 1 (CL1) are observed to insert into the translocation pathway from opposite sides (Fig. 2A), suggesting potential roles in VIAAT function. LL4 is positioned in close proximity to TM1b, placing it near the central transport pathway. The highly charged residue Asp364 in LL4 forms salt bridges with Lys296 and Lys298 in LL3, which connects TM5 and TM6a. LL4 also interacts with TM1b through polar contacts between Glu368 and Tyr139, as well as between the backbone carbonyl oxygens of Glu368 and Leu135. In addition, Ile370 in LL4 engage in hydrophobic interactions with Leu135, Phe354, Val281, and Ala358 (Fig. 2E). To investigate the functional role of LL4, we replaced LL4 (residues 363-371) with a (GS)_n_ linker (VIAAT-GS) and first assessed its thermostability. Compared with WT VIAAT, VIAAT-GS exhibited markedly reduced thermostability (SI Appendix, Fig. S1C). Furthermore, mIPSC amplitudes in neurons expressing VIAAT-GS were almost completely abolished (Fig. 2D). These results indicate that LL4 is critical for VIAAT stability and function. We propose that the LL4 may move toward the translocation pathway and occlude the luminal vestibule through interactions with surrounding TMs during the transition from a luminal-open to a luminal-closed conformation.

The large CL1, composed of two β-strands and a short α-helix, wedges between TM2, TM3, TM6b, and TM10. Arg181 and Tyr171 engage TM6b via hydrogen bonds with the carboxylate oxygen of Pro324 and the side chain of Glu327, respectively. In addition, Tyr186 inserts into a hydrophobic pocket formed by Leu166, Val206, Phe476, and Leu468 (Fig. 2F, G). These intensive interactions likely contribute to maintaining the overall conformation of VIAAT. To validate our hypothesis, we simultaneously substituted Tyr171, Arg181 and Tyr186 with alanines (Y171A/R181A/Y186A) and measured mIPSCs. Compared with WT VIAAT, the Y171A/R181A/Y186A mutant exhibited markedly reduced mIPSC amplitudes, indicating that CL1 is critical for VIAAT function (Fig. 2D).

### GABA binding site of VIAAT

To elucidate the molecular basis of substrate recognition by VIAAT, we determined the structure of VIAAT-GABA. Residues from TM1, TM3, and TM6 together form the central binding pocket in which GABA binds. The helical arrangement within this pocket is nearly identical to that of VIAAT-apo, with a root-mean-square deviation (r.m.s.d.) of approximately 0.4 Å. However, in the GABA-bound structure, the luminal access pathway is occluded by Phe133 and Phe315. Together with the cytosolic gate residue Phe322, these interactions block the substrate translocation pathway from both sides, indicating that VIAAT-GABA adopts an occluded state (Fig. 3A).

**Fig. 3.**
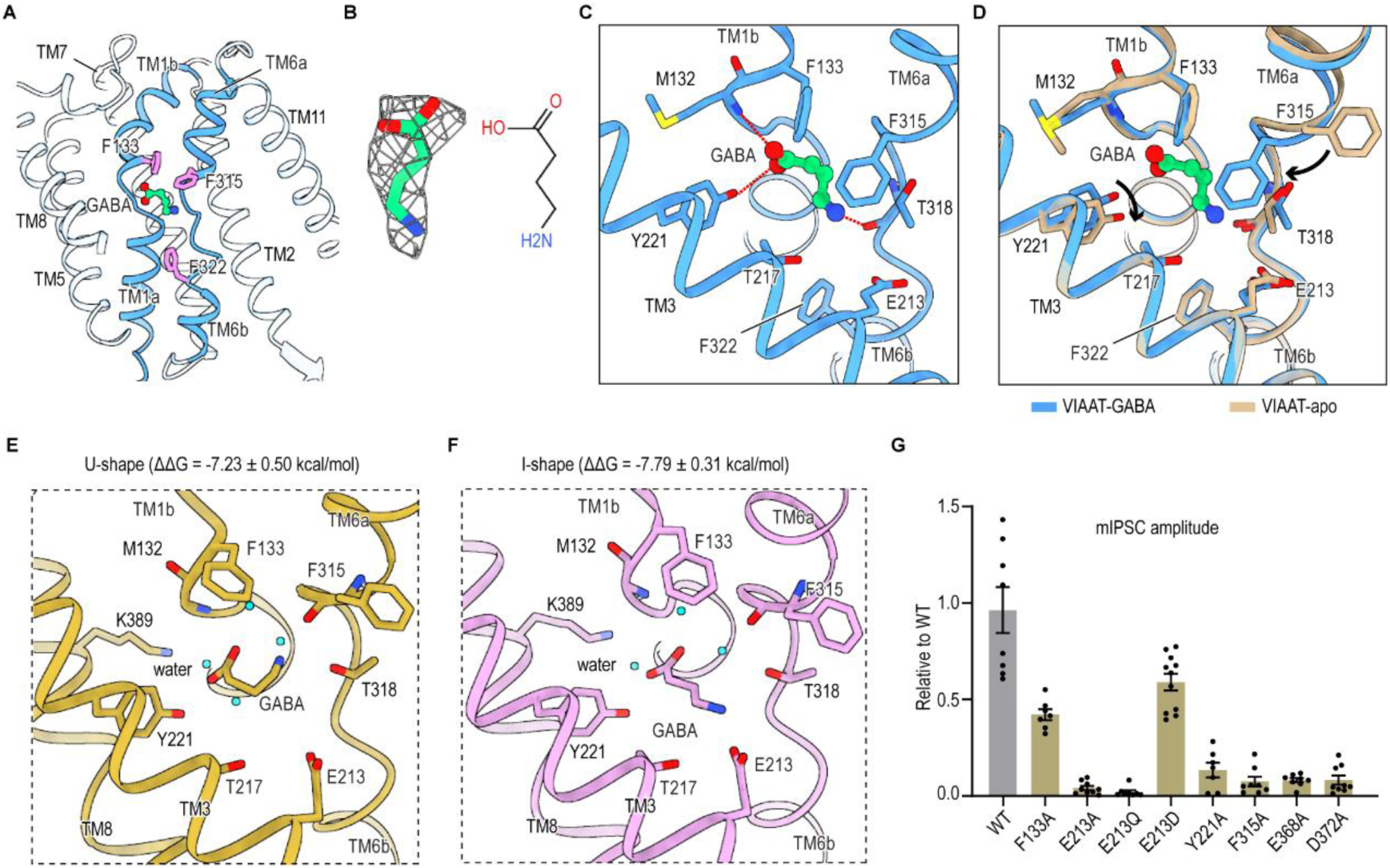
GABA binding and recognition. **(A)** The GABA-bound structure is locked in a luminal-occluded state by Phe133, Phe315 and Phe322. **(B)** Left, GABA is overlaid with the corresponding cryo-EM density (mesh). Right, chemical structure of GABA. **(C)** GABA in the central binding pocket of VIAAT-GABA. Hydrogen bonds are indicated by red dashed lines. **(D)** Superimposition of VIAAT-apo (wheat) and VIAAT-GABA (blue). GABA is shown as a green ball-and-stick model. Black arrows indicate the conformational rearrangements of Tyr221 and Phe315. **(E-F)** Molecular dynamics simulations reveal two binding modes of GABA in VIAAT: the U-shape (**E**) and the I-shape (**F**). **(G)** mIPSCs recorded from hippocampal neurons after knockdown of endogenous VIAAT and rescue with WT human VIAAT or its mutants. mIPSC amplitudes were normalized to those of VIAAT-WT (n = 7-11 cells per condition). Data are shown as mean ± s.e.m.

In the central pocket, a prominent elongated density was observed, matching the size and shape of GABA and accommodating its carboxylate and amino moieties (Fig. 3B). The position and orientation of GABA are analogous to those observed in structures of the GABA transporter (GAT1)(30) and taurine transporter (TauT)(26), both of which are capable of transporting GABA (SI Appendix, Fig. S5). GABA is enclosed by four aromatic residues Phe133, Tyr221, Phe315, and Phe322 that likely engage in polar-π interactions. The two carboxylate oxygens of GABA are oriented toward Phe133 and Tyr221, whereas the amine group is positioned between Phe315 and Phe322. Additionally, the carboxylate group of GABA forms potential hydrogen bonds with the hydroxyl group of Tyr221 and the backbone amide of Met132, while the amine group forms a hydrogen bond with the backbone carbonyl of Thr318 (Fig. 3C). Thus, Phe322 not only contributes to the cytosolic gating but also directly participates in substrate recognition within the central binding pocket.

To assess the functional importance of these interactions, we substituted Phe133, Tyr221, and Phe315 with alanine and measured their effects on inhibitory synaptic transmission. The Tyr221Ala, Phe315Ala and Phe322Ala mutants caused marked reductions in mIPSC amplitudes, whereas Phe133Ala produced a modest decrease (Fig. 3G). Together, these results indicate that the aromatic residues surrounding the substrate-binding pocket are critical for GABA recognition and synaptic transmission.

Comparison of VIAAT-apo and VIAAT-GABA structures reveals that most substrate-coordinating residues adopt similar conformations, except for Tyr221 and Phe315 (Fig. 3D). Tyr221 exhibits two alternative side chain conformations, whereas Phe315 undergoes a pronounced displacement. In the VIAAT-apo structure, the aromatic side chain of Phe315 swings away from the central pocket, opening the luminal translocation pathway. In contrast, in the VIAAT-GABA structure, Phe315 rotates back toward the substrate, thereby occluding the luminal pathway. Together with the alternative conformations of Tyr221, this rearrangement further restricts solvent access to the central binding pocket from the luminal side (Fig. 3D). These observations indicate that Phe315 functions as a luminal gate in addition to its role in substrate binding. This is reminiscent of observations in the choline transporter CHT1 (SLC5A7)(31), in which a conformational rearrangement of a single residue blocks the intracellular solvent pathway.

To explore the molecular determinants of substrate specificity among VIAAT, GAT1, and TauT, we superimposed their central binding pockets (SI Appendix, Fig. S5). This analysis revealed that most surrounding residues differ among the three transporters, with the exception of two conserved positions corresponding to Tyr221 and Phe315 in VIAAT. Tyr221 aligns with Tyr140 in GAT1 and Tyr138 in TauT, residues previously reported to be indispensable for substrate binding in those transporters, supporting our model that Tyr221 plays a prominent role in substrate recognition in VIAAT. Phe315 corresponds to Phe294 in GAT1 and Phe300 in TauT, both of which have been identified as extracellular gating residues. This conservation is consistent with our observation that conformational rearrangements of Phe315 are critical for opening and closing the luminal translocation pathway in VIAAT.

Additionally, we performed molecular dynamics (MD) simulations to investigate the stability of GABA binding to VIAAT (Fig. 3E, F and SI Appendix, Fig. S6). The simulations revealed that GABA binds in two distinct modes: a U-shape and an I-shape. Across all simulations, the I-shape accounted for approximately 58%, and the U-shape for 36% (with the remaining 6% representing transitional states), indicating the flexible binding of GABA in VIAAT. In the U-shape mode, GABA forms an intramolecular hydrogen bond between its amino and carboxyl groups, which prevents the amino group from interacting with Glu213. In the I-shape mode, the amino group forms a salt bridge with Glu213, while the carboxyl group engages in a moderate electrostatic interaction with Lys389. Tyr221 and Met132 interact with GABA via hydrogen bonds in both binding modes. Notably, the I-shape mode closely resembles the binding mode observed in our cryo-EM structure. The calculated binding free energies for the two modes were similar: -7.23 kcal/mol for the U-shape and -7.97 kcal/mol for the I-shape. We further calculated the binding free energies to assess the contribution of key pocket residues (SI Appendix, Fig. S6B, C). In the I-shape mode, Glu213 contributed -0.58 kcal/mol and Tyr221 contributed -2.84 kcal/mol to GABA binding. In the U-shape mode, Glu213 made no appreciable contribution, consistent with the intramolecular hydrogen bond that sequesters its amino group from the pocket, whereas Tyr221 contributed a similar value. Together, these simulations demonstrate that GABA adopts two binding poses within the VIAAT pocket, with the I-shape mode being both energetically favorable and structurally consistent with our experimental data.

### Transport mechanism of VIAAT

Transporters with LeuT-like fold follow a rocker-switch mechanism, cycling between cytosolic-facing and periplasmic/luminal-facing conformations to alternately expose the substrate-binding site to opposite sides of the membrane and drive substrate transport. drSLC38A9 shows close structural similarity to VIAAT and has been solved in a cytosolic-facing state. To gain insight into the transport cycle of VIAAT, we therefore generated a cytosolic-facing model of VIAAT using the drSLC38A9 structure as a template.

Comparison of VIAAT-apo with this cytosolic-facing model revealed pronounced conformational changes on both the luminal and cytosolic sides (Fig. 4A-C). On the luminal side, the luminal segments of TM10 and TM3 undergo substantial inward rotations, while TM1b and TM6a show modest displacements (Fig. 4B). Consistent with our hypothesis, LL4 moves closer to the center of the luminal permeation pathway (Fig. 4B, D-E). The inward rotations of TM1b, TM3, TM6a, and TM10, along with the repositioning of LL4, enable specific interactions that seal the luminal gate. In this model, Asp372 in LL4 forms salt bridges with Asn227 and Asn231, whereas Glu368 in LL4 forms a hydrogen bond with the main-chain amide of Phe453 in TM10. The side chain of Phe453 further projects into the central pathway and engages in hydrophobic interactions with Phe133 in TM1b, Leu220 in TM3 and Met457 in TM10, thereby contributing to closure of the luminal gate (Fig. 4E). On the cytosolic side, in contrast to the pronounced movement of the luminal segments of TM3 and TM10, their cytosolic segments remain largely unchanged (Fig. 4C). The most prominent conformational changes involve outward movements of TM4, TM5, TM8, and TM9, accompanied by TM1a and the N-terminal loop shifting toward the membrane plane. These rearrangements disrupt the hydrophobic interactions mediated by Phe322 and surrounding residues that occlude the cytosolic gate in the luminal-facing state (Fig. 4F). In particular, the large displacements of TM5 and TM8 create space for substrate access between the central pocket and the cytosol, thereby opening the cytosolic gate. Notably, mutation of Val263, located in the cytosolic half of TM5, has been linked to genetic epilepsy, highlighting the functional importance of this region in VIAAT transport(10). By contrast, TM2, TM7, and TM11 undergo minimal rearrangement during this transition, suggesting that they constitute a relatively rigid scaffold domain.

**Fig. 4.**
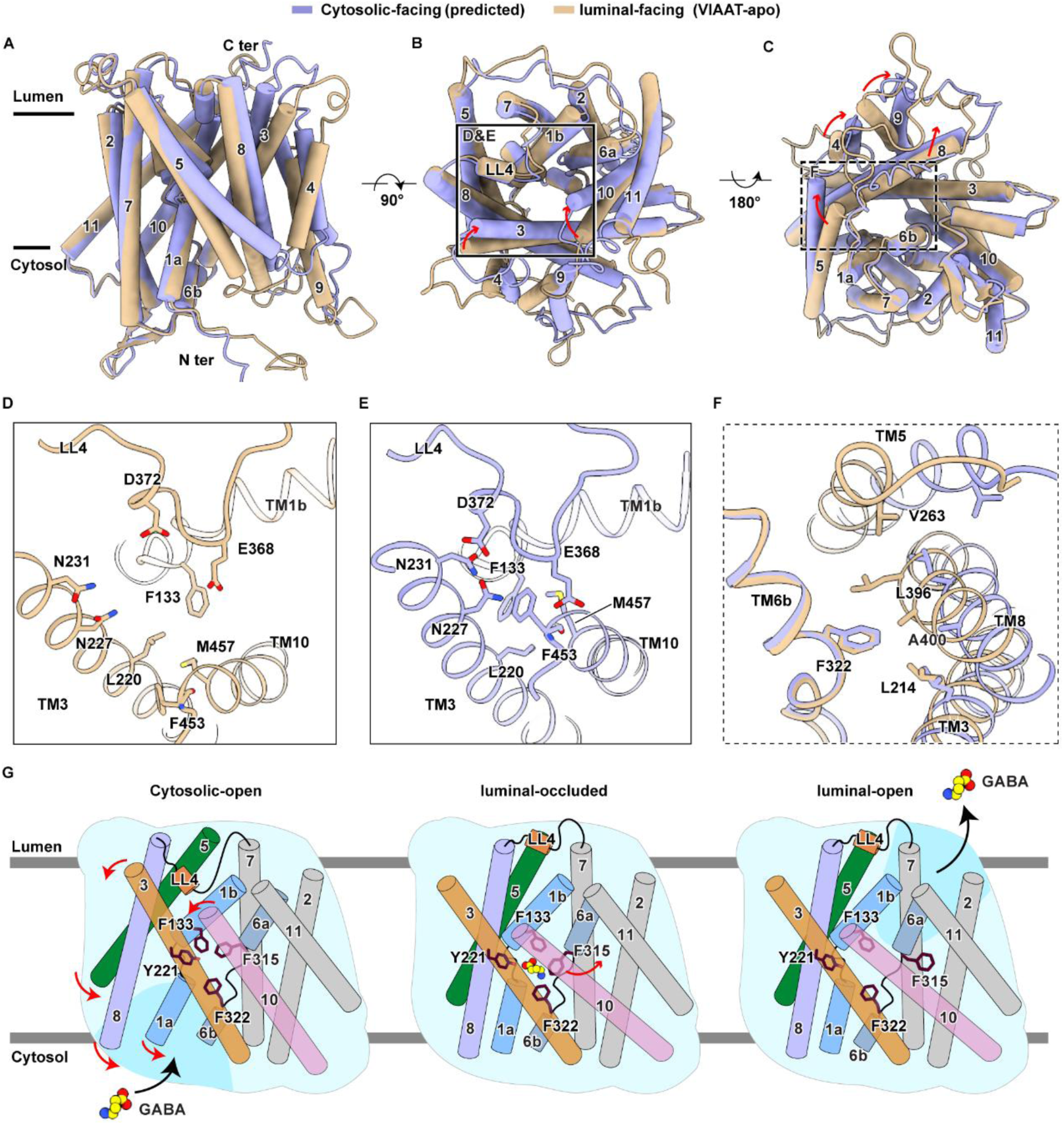
Mechanism of VIAAT-mediated GABA transport. **(A-C)** Superimposition of VIAAT-apo (wheat) and the predicted cytosolic-facing model (purple), viewed along the membrane plane (**A**), from the luminal side (**B**) and from the cytosolic side (**C**). Red arrows indicate pronounced conformational rearrangements. **(D-E)** Enlarged views of the boxed region in (**B**), showing the conformational changes on the luminal sides between VIAAT-apo (**D**) and in the cytosolic-facing model (**E**). **(F)** Enlarged view of the boxed region in **c**, showing structural transitions in the cytosolic sides. **(G)** Schematic of VIAAT-mediated GABA transport. GABA-binding and recognition residues are shown. Red arrows indicate major conformational changes.

As with other vesicular neurotransmitter transporters, VIAAT transport activity is driven by the proton electrochemical gradient. Following substrate release, luminal proton binding is required for VIAAT to return to a cytoplasm-facing state. However, no specific residue has yet been identified that directly senses protons or undergoes protonation during the transport cycle. We observed a single acidic residue, Glu213, within the central binding pocket, with one of its carboxylate oxygens positioned approximately 4.8 Å from the amine group of GABA (Fig. 3C). To investigate the functional importance of Glu213, we generated alanine (E213A), glutamine (E213Q), and aspartate (E213D) substitutions. Previous studies reconstituting VIAAT into liposomes showed that the E213A mutation abolished uptake activity(32), and we confirmed that it severely impaired mIPSCs (Fig. 3G). The neutral substitution (E213Q) also eliminated transport function, whereas the conservative substitution (E213D) retained most activity. These results indicate that a negative charge at this position is critical and that the side chain volume of Glu213 also contributes to its optimal function. Together, these findings suggest that Glu213 plays a critical functional role, potentially by participating in proton-coupled transport or by directly interacting with the substrate. Consistent this interpretation, in both VMAT2 and VAChT, residues implicated in proton coupling also contribute to substrate binding(15–17). Additionally, we identified two acidic residues Asp372 and Glu368 in the luminal side of VIAAT (Fig. 4D). Alanine substitutions of either residue Asp372 or Glu368 completely abolished transport activity, underscoring their essential roles (Fig. 3G). Given their acidic nature and location within LL4, we speculate that Asp372 and Glu368 may contribute to LL4-mediated gating or proton coupling during the transport cycle.

## Discussion

In this study, we captured the VIAAT structure in the apo luminal-open state and the GABA-bound occluded state. Together with a predicted cytosolic-open state and functional validation, we propose that VIAAT transports GABA into synaptic vesicles via a rocker-switch mechanism (Fig. 4G). In the cytosolic-open state, GABA enters the binding pocket and is recognized by a conserved aromatic cage comprising Phe133, Tyr221, Phe315, and Phe322. Substrate binding then promotes transition to an occluded state, in which access from both sides of the membrane is restricted. This transition is accompanied by inward movements of TM1a, TM5 and TM8, which establish hydrophobic interactions that seal the cytosolic gate. Concurrently, outward rotations of TM3, TM6a and TM10, together with movement of LL4 away from the permeation pathway, open the luminal vestibule. Subsequent rotation of Phe315 away from the substrate further opens the luminal pathway, allowing GABA release into the vesicle lumen. Notably, TM2, TM7 and TM11 remain largely unchanged during these conformational transitions, suggesting that they form a relatively rigid scaffold throughout the transport cycle.

It remains controversial whether VIAAT transport activity depends on chloride in addition to the proton electrochemical gradient(32). In the GABA-bound structure, but not in VIAAT-apo, we observed an additional density adjacent to the substrate (SI Appendix, Fig. S7A), reminiscent of the Na1 site in members of the SLC6 superfamily, where a bound sodium ion is essential for transport. This density is positioned to be coordinated by the carboxylate of GABA, the hydroxyl group of Tyr221, and the side chains of Asn127 and Thr217. Moreover, mutation of Asn127 to alanine markedly impaired synaptic transmission, suggesting that Asn127 plays an important functional role, potentially by coordinating this density (SI Appendix, Fig. S7B). Definitive assignment of this feature, and clarification of whether chloride contributes to VIAAT transport, will require robust cell-or liposome-based uptake assays, which were not available in the present study.

Given that VIAAT transports both GABA and glycine, albeit with lower affinity for glycine, we sought to determine the structure of VIAAT in complex with glycine. However, due to the smaller size of glycine compared with GABA and the limited resolution of the reconstruction, we could not confidently assign its position in the binding pocket. Further studies are required to define the molecular basis of glycine recognition by VIAAT.

Overall, our work provides structural and functional insight into how VIAAT recognizes GABA and mediates its uptake into synaptic vesicles. These findings establish a mechanistic framework for understanding vesicular inhibitory neurotransmitter loading and help explain how epilepsy-associated VIAAT variants disrupt transporter function.

## Methods

### Cloning, expression and purification of human VIAAT

The full-length human VIAAT gene (*SLC32A1*, UniProt ID: Q9H598) was synthesized and cloned into the pEGBacMam vector containing a 3C protease cleavage site and a Twin-Strep affinity tag at its C-terminus. Baculovirus was produced in Sf9 cells using the Bac-to-Bac system and P2 viruses were used to transfect HEK293S GnTI^−^ cells. HEK293S GnTI^−^ cells were cultured in Freestyle 293 Expression Medium (Gibco) supplemented with fetal bovine serum (FBS) (1% v/v) in a 37 ℃ incubator with 5% CO_2_. When the cell density reached approximately 3.0 × 10^6^ cells/mL, P2 viruses were added to the culture at a 1:20 (v/v) ratio. After 10 hours, 10 mM sodium butyrate was added to induce protein expression, and the cells were cultured at 30 ℃ for another 60 hours before being harvested.

The cell pellets were resuspended in lysis buffer (20 mM Tris pH 8.0, 150 mM NaCl, 1% (w/v) glyco-diosgenin (GDN, Anatrace), 0.8 µM aprotinin, 2 µg/mL leupeptin, 2 µM pepstatin A and 1 mM phenylmethylsulfonyl fluoride) supplemented with 5% glycerol, and lysed at 4 ℃ for 2 hours. Following centrifugation at 4 ℃ and 186,009 g for 1 hour, the supernatant was filtered through a 0.45 µm polystyrene membrane. The filtrate was then incubated with Strep-Tactin resin at 4 ℃ for 1 hour. The resin was pre-equilibrated with the buffer A (20 mM Tris pH 8.0, 150 mM NaCl, 0.005% (w/v) lauryl maltose neopentyl glycol (LMNG, Anatrace) and 5% glycerol). Then the resin was washed with 30 column volumes of buffer A. The protein was eluted by buffer A supplemented with 5 mM desthiobiotin and subsequently concentrated to 1mL. Purified protein was further processed by size-exclusion chromatography on a Superose 6 10/300 GL column (GE Healthcare) in buffer A without 5% glycerol. The protein peak fractions were collected and concentrated for cryo-EM analysis. For the VIAAT-GABA complex, the protein was incubated with 100 mM GABA (Sigma-Aldrich) at 4 ℃ for 30 min prior to cryo-EM sample preparation.

### Electrophysiology of cultured neurons

Dissociated hippocampal neuron cultures were prepared from wild-type C57BL/6 mouse pups at postnatal day 0 (P0). Neurons were seeded onto poly-L-lysine-coated glass coverslips and maintained in Neurobasal-A medium supplemented with 2% B-27 and 1% GlutaMAX at 37 °C in a 5% CO₂ incubator. On day in vitro 3 (DIV3), cultures were transduced with a lentivirus encoding an shRNA (U6-shRNA-CMV-GFP) targeting the VIAAT mRNA sequence 5 ′ - CCAAGTTCAGTCTGCTGTGTA-3′ to knock down endogenous VIAAT. On DIV5, cells were transduced with lentiviral constructs expressing either wild-type or mutant VIAAT with an IRES-mScarlet tag for rescue experiments. To prevent shRNA-mediated silencing of the rescue VIAAT, the target sequence (5 ′ -CCAAGTTCAGTCTGCTGTGCA-3 ′) in the complementary DNA (cDNA) was replaced with a synonymous mutant sequence (5 ′ - CTAAATTTAGCCTTCTTTGTA-3 ′) in all rescue plasmids. Except for Lenti-U6-shRNA-CMV-GFP, which was purchased from VectorBuild, lentiviruses used for VIAAT rescue were packaged in HEK293T cells. Specifically, HEK293T cells were co-transfected with the corresponding lentiviral transfer plasmids, psPAX2 and pMD2.G using PEI. After 48 hours, the supernatant was collected, filtered through a 0.45 µm syringe filter, and stored at -80 °C for subsequent use.

Whole cell voltage clamp recordings were performed on neurons between DIV14 and DIV16. The pipette solution contained 40 mM CsCl, 90 mM K-gluconate, 2 mM MgCl_2_, 3.5 mM KCl, 0.1 mM EGTA, 2 mM Mg-ATP, 1 mM Na_2_-GTP, 10 mM phosphocreatine, 5 mM QX-314, and 10 mM HEPES (pH 7.4, 300 mOsm). The bath solution contained 140 mM NaCl, 5 mM KCl, 2 mM CaCl_2_, 2 mM MgCl_2_, 10 mM glucose, 0.05 mM D-AP5, 0.02 mM CNQX, and 10 mM HEPES (pH 7.4, 310 mOsm). Neurons were clamped at −70 mV and series resistance was compensated to approximately 5 MΩ. Electrical stimulation was applied using a bipolar electrode. Signals were acquired using a Multiclamp 700B amplifier and a Digidata 1550B digitizer, and data were analyzed with pClamp10.

### Western blotting

Hippocampal and cortical neurons were cultured from C57BL/6 mouse embryos at embryonic day 17 (E17). Cells were plated on poly-D-lysine-coated 24-well plates at a density of 250,000 cells per well and maintained in Neurobasal medium supplemented with 2% B-27, 0.25% GlutaMAX, and 1% penicillin–streptomycin. On DIV3, cultures were transduced with lentivirus (U6-shRNA-CMV-GFP) and harvested on DIV7-10. Cells were lysed in buffer (25 mM Tris-HCl pH 8.0, 150 mM NaCl, 10% glycerol, 0.8 µM aprotinin, 2 µg mL⁻¹ leupeptin, 2 µM pepstatin A, and 1% NP-40) at 4 °C for 15 min. Lysates were cleared by centrifugation at 4 °C and 15,000g for 10 min, and equal amounts of total protein were separated by 4-15% SDS-PAGE. Proteins were transferred to nitrocellulose membranes and immunoblotted with primary antibodies against VIAAT (mouse, 1:1,000, Synaptic Systems, 131011) and GAPDH (mouse, 1:50,000, Proteintech, 60004-1-Ig), followed by IRDye 800CW-conjugated secondary antibodies (goat anti-mouse IgG, 1:10,000, LI-COR). Protein bands were visualized using an Odyssey DLx imaging system (LI-COR).

### RNA isolation and RT-qPCR

Neurons were prepared as described above for western blotting. Total RNA was extracted using TRIzol reagent, and equal amounts of RNA from each sample were reverse-transcribed into complementary DNA (cDNA) using HiScript IV RT SuperMix (Vazyme, R423-01) according to the manufacturer’s instructions. The synthesized cDNA was diluted and used for real-time PCR with ChamQ Blue Universal SYBR qPCR Master Mix (Vazyme, Q312-02) on a 7500 Real-Time PCR system (Applied Biosystems). Ct values were collected, and the ΔΔCt method was used to calculate normalized fold changes in relative mRNA expression.

### Cryo-EM sample preparation and data collection

A 4 µL aliquot of each protein sample (2.5 mg/mL) was applied to glow-discharged Quantifoil R2/1 200 mesh copper holey carbon grids. The grids were then blotted for 3 s at 8 ℃ under conditions of 95% humidity, followed by plunge freezing into liquid ethane cooled by liquid nitrogen, performed in the Vitrobot Mark IV. Cryo-EM data were collected on a 300-kV Titan Krios microscope (Thermo Fisher Scientific) using a K3 direct electron detector (Gatan) and an energy filter. Video stacks were automatically collected using EPU software at a calibrated magnification of 81,000× in the super-resolution mode. The pixel size of micrographs was 0.5275 Å with a defocus range of -1.0 to -2.0 μm. Each micrograph was recorded with a total dose of 50 e-/Å^2^ at a dose rate of ∼ 25 e-/pixel/s.

### Cryo-EM data processing

All Cryo-EM data were processed using cryoSPARC(33). For the VIAAT-apo dataset, a total of 5,351 movies were collected. Beam-induced motion was corrected using patch motion correction, and the contrast transfer function (CTF) was estimated using patch CTF estimation. Micrographs with a CTF fit resolution exceeding 4 Å were discarded, resulting in 4,975 micrographs for further processing. A total of 4,979,976 particles were initially picked using the blob picker and extracted with a box size of 256 pixels. Junk particles were removed through two rounds of 2D classification. Classes exhibiting elliptical micelle shapes were selected for ab-initio reconstruction to generate an initial model. This initial model was used as a good reference for heterogeneous refinement, whereas junk classes from the initial ab-initio reconstruction were used as additional references to remove non-target particles. Additionally, initial junk particles from the first round of 2D classification were subjected to ab-initio reconstruction to generate poor references for heterogeneous refinement. After multiple rounds of multi-reference heterogeneous refinement, 197,252 good particles were obtained. Ab-initio reconstruction was then performed to further remove poor quality particles. The resulting good particle class was refined to a helical distinct map at 4.1 Å resolution using non-uniform refinement. Without-alignment 3D classification was used to remove low-resolution particles, yielding 37,695 particles, which were then reconstructed to 3.7 Å using non-uniform refinement followed by local refinement. These particles were then subjected to 2D classification, and the helical distinct classes were selected for Topaz training. A total of 4,710,296 particles were selected by Topaz extraction. Subsequently, a seed-facilitated classification strategy was used to enrich good particles from the particle batch selected by Topaz(34), ultimately generating 334,879 good particles. Two rounds of without-alignment 3D classification were then performed to remove poor-quality and low-resolution particles, followed by non-uniform refinement and local refinement, yielding a final map at 3.2 Å resolution from 34,411 particles. For the VIAAT-GABA dataset, a total of 5,929 movies were collected. The initial particle batch was selected using the template picker, and the processing strategy was identical to that used for the VIAAT-apo dataset.

### Model building

The AlphaFold3-predicted model of VIAAT was roughly fitted into the cryo-EM density map of VIAAT-apo by UCSF ChimeraX(35). The fitted model was manually adjusted and corrected in Coot(36) and subsequently refined in real space using Phenix(37). The resulting VIAAT-apo model served as the initial template for generating and refining models for VIAAT-GABA. Geometry constraints for GABA were produced by elBOW tool in Phenix(37). The model validation was performed using MolProbity(38) and Phenix(37), with refinement and validation statistics summarized in the SI Appendix, Table 1. All figures were prepared using UCSF ChimeraX(35).

### FSEC-based thermostability assay

Constructs encoding wild-type or mutant VIAAT with a C-terminal YFP tag were used. Plasmids were transiently transfected into HEK293S GnTI⁻ cells. At 60 hours post-transfection, cells were solubilized in lysis buffer (20 mM Tris-HCl pH 8.0, 150 mM NaCl, 1% (w/v) GDN, and protease inhibitors) at 4 °C for 2 hours. Cell debris was removed by centrifugation at 40,000g for 20 min, and the resulting supernatants were heated at various temperatures for 10 min. Denatured proteins were removed by ultracentrifugation at 40,000g for 20 min. VIAAT proteins were separated using a Superose 6 Increase 10/300 GL column for fluorescence size-exclusion chromatography (FSEC), and YFP fluorescence was measured (excitation 515 nm, emission 535 nm). Peak heights corresponding to VIAAT were normalized to those from untreated samples (no heat treatment). Melting temperatures (Tₘ) were determined by fitting the normalized data to a four-parameter dose–response equation in GraphPad Prism 9.

### Molecular Dynamics Simulations

The Cryo-EM structure of VIAAT-GABA was employed as the initial structure for the MD simulations. Here, only the residues 109-524 have been considered, and the N-terminus was ignored due to the unobservable density map and low pLDDT values (<50) in AlphaFold3(39) predictions. During system preparation, the pKa values of acidic and basic residues were predicted using the PROPKA3 program(40). All acidic residues’ pKa values were lower than the buffer pH 8.0, and basic residues’ pKa values were higher than 8.0, indicating that the side chains of the acidic residues were deprotonated and those of the basic residues were protonated. The GABA ligand was modeled as a zwitterion with a neutral net charge. The membrane system was constructed using the CHARMM-GUI web server(41). The termini of the protein were capped by neutral groups, namely the acetyl (ACE) N-terminal cap and methylamine (CT3) C-terminal cap. The GABA was described by the CHARMM General Force Field (CGenFF)(42) provided in CHARMM-GUI. The position of the protein in the membrane was calculated with the Orientation of Protein in Membranes (OPM) PPM 2.0 web server(43). The lipid bilayer was composed of POPC and POPE with a ratio of 3:1, and in total, 158 POPC and 52 POPE were added to the system. About 21500 TIP3P waters were added on top and bottom of the bilayer system, and later, the model was neutralized with 150 mM NaCl. The system size was 95×95×118 Å^3^, and the periodic boundary conditions were established based on this box size. Eventually, the AMBER simulation input files were generated for further simulations(44). The CHARMM36m forcefield(45) was used to describe the protein behavior, and the CHARMM all-atom additive forcefield(46) for lipids.

First of all, the systems were minimized with position restraints on protein and ligand (10 kcal/mol/Å^2^), and lipids (2.5kcal/mol/Å^2^). Following the minimization, a 250 ps heating process and five sequential equilibrium steps were carried out with gradually reducing the position restraints on lipid, protein backbone, and protein side chains. Finally, three replicates of 300-ns production MD simulations were performed under the NPT ensemble. In the simulations, the semi-isotropic pressure of the system was controlled by the Monte Carlo barostat(47), and the temperature was maintained by the Langevin thermostat algorithm with a collision frequency of 1.0 ps^-1^. The nonbonding interaction cutoff value was set to 12 Å, with a switching function starting from 10 Å for the van Der Waals interactions. The particle-mesh Ewald (PME) method was used to describe the long-range electrostatic interaction. The SHAKE algorithm was set to constrained bonds involving hydrogen. All the simulations were done with AMBER24, and the binding free energy was calculated by the MM-GBSA method(48) using the last 100 ns trajectories.

### Animal use statement

All mouse experimental protocols were approved by the Institutional Animal Care and Use Committee (IACUC) of the Interdisciplinary Research Center on Biology and Chemistry (IRCBC), Shanghai Institute of Organic Chemistry, Chinese Academy of Sciences, under approval number ECSIOC_2023-14.

## Data availability

The 3D cryo-EM density maps and coordinates of VIAAT-apo, VIAAT-GABA have been deposited in the Electron Microscopy Data Bank (EMDB) under the accession numbers EMD-81415 and EMD-81416 and in the Protein Data Bank (PDB) under accession codes 27TQ and 27TR, respectively.

## Acknowledgments

We thank the Cryo-Electron Microscopy Center at the Interdisciplinary Research Center on Biology and Chemistry, Shanghai Institute of Organic Chemistry for the help with data collection. This work was supported by the Shanghai Basic Research Pioneer Project (to J.Y.), Shanghai Key Laboratory of Aging Studies (19DZ2260400, to J.Y.), startup grant of ShanghaiTech University (to J. G.) and Shanghai Pujiang Program (to J. G.).

## Author contributions

J.Y. and J.G. conceived and supervised the project. J.C. performed protein purification, cryo-EM sample preparation, electrophysiology, and thermostability assays. N.C. performed molecular dynamics (MD) simulations. J.C. and J.Y. performed cryo-EM data collection, model building, and data analysis. J.Y. and J.C. prepared the figures. J.Y., J.C., and J.G. wrote the manuscript.

## Materials & Correspondence

Correspondence and requests for materials should be addressed to Jie Yu.

## Supporting Information

### Supplementary Figure and Figure Legends

**Fig. S1.**
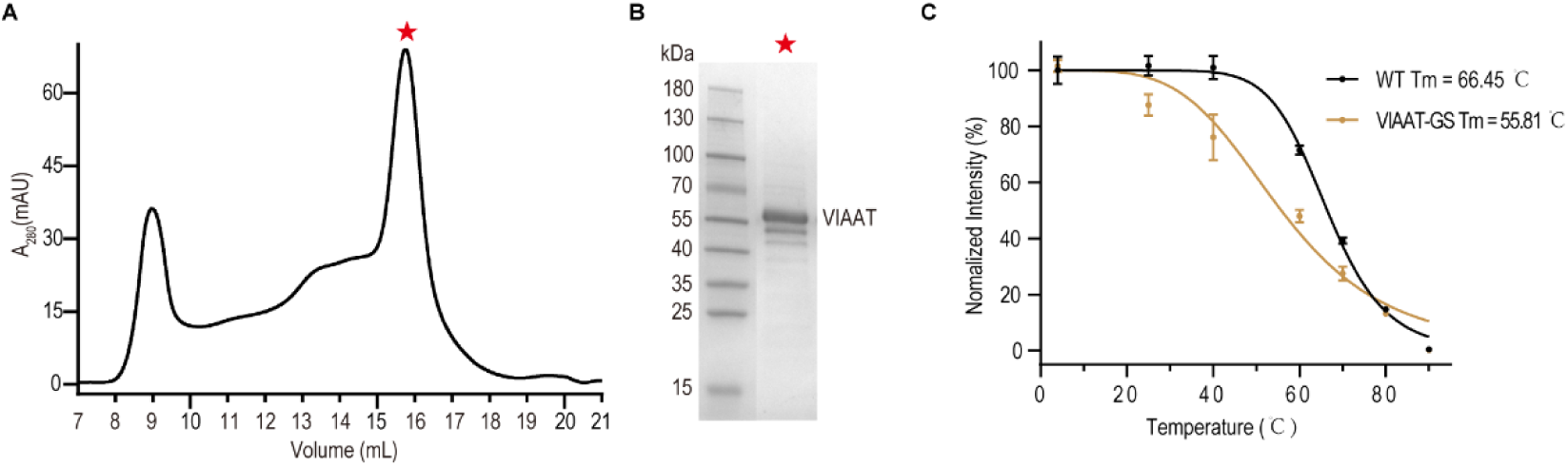
Biochemical characterization of the hVIAAT. **(A)** Size-exclusion chromatogram (Superose 6 10/300 GL column) profile of the purified VIAAT protein. The red star indicates peak fractions of purified VIAAT collected for cryo-EM sample preparation. **(B)** Coomassie-blue-stained SDS-PAGE gel of purified VIAAT protein. **(C)** Thermal stability assays of WT VIAAT and VIAAT-GS. Data are presented as mean ± SEM, n = 3 biologically independent samples.

**Fig. S2.**
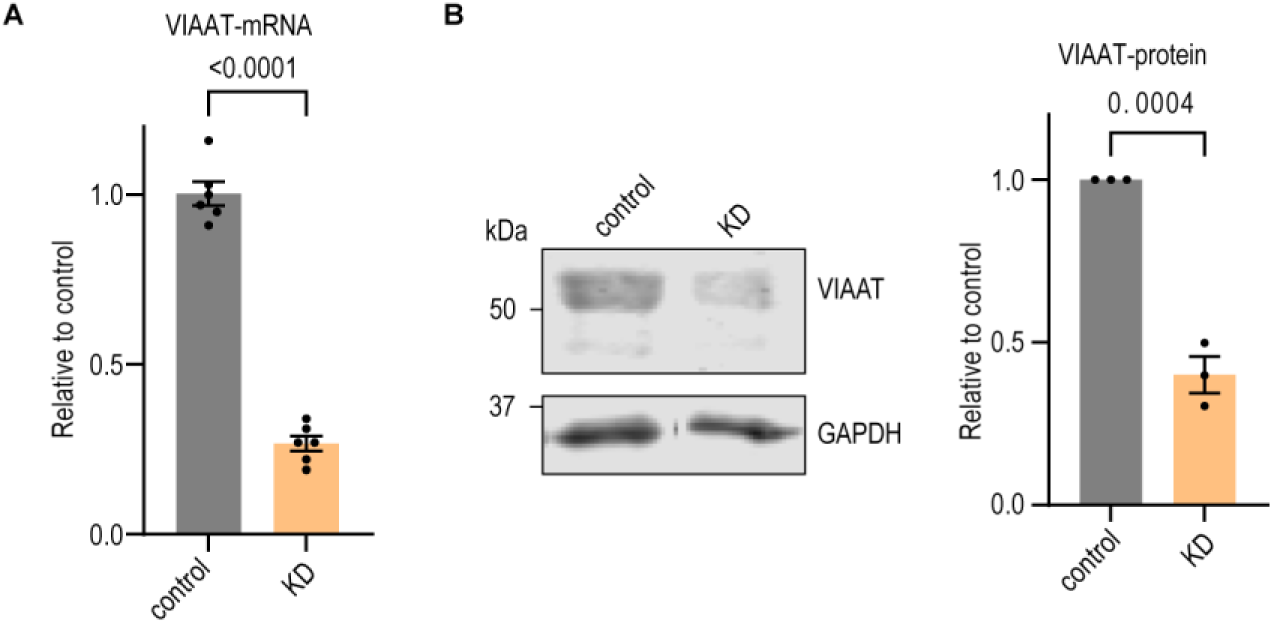
Validation of endogenous VIAAT knockdown (KD) by shRNA. **(A)** Relative VIAAT mRNA levels in control and KD samples, measured by qPCR. Data are shown as mean ± s.e.m. from six independent experiments. Two-sided Student’s t-test; P values are indicated in the figure. **(B)** Western blot analysis of endogenous VIAAT protein levels after knockdown. Bar graph shows mean ± s.e.m. from three independent experiments. Two-sided Student’s t-test; P values are indicated in the figure.

**Fig. S3.**
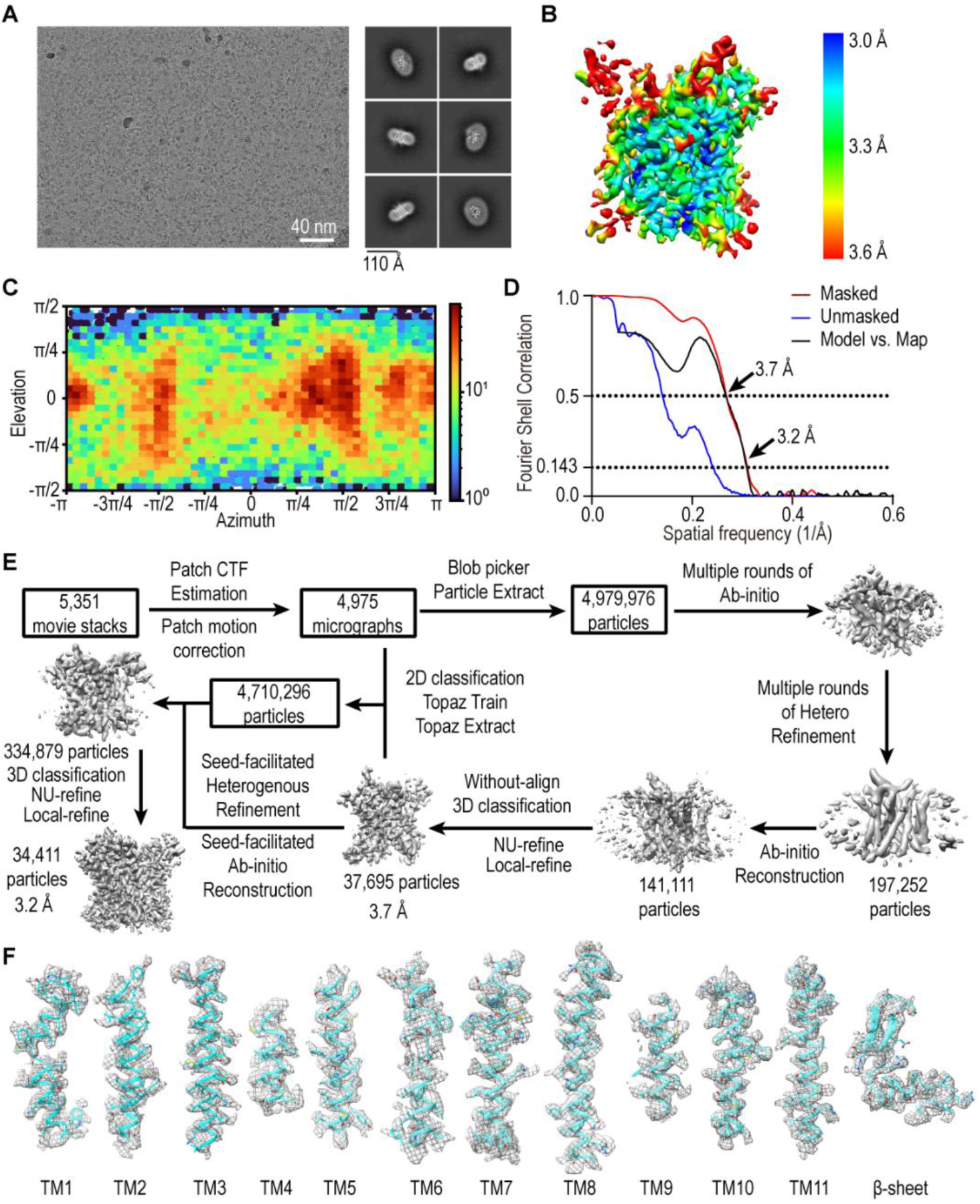
Cryo-EM reconstruction of VIAAT-apo. **(A)** Representative micrograph (left) (scale bar, 40 nm) and 2D class averages (right). **(B)** Local resolution distribution of VIAAT-apo. **(C)** Angular distribution of the final reconstruction. **(D)** Gold standard Fourier Shell Correlations (FSC) curves of unmasked map (blue), masked map (red) and model versus map (black), marked with resolutions corresponding to FSC = 0.5 and 0.143. **(E)** Flow chart for cryo-EM data processing of VIAAT-apo. Further details can be found in the Methods. **(F)** Cryo-EM densities of TM1-11 and the β-sheet in VIAAT-apo, contoured at 0.3.

**Fig. S4.**
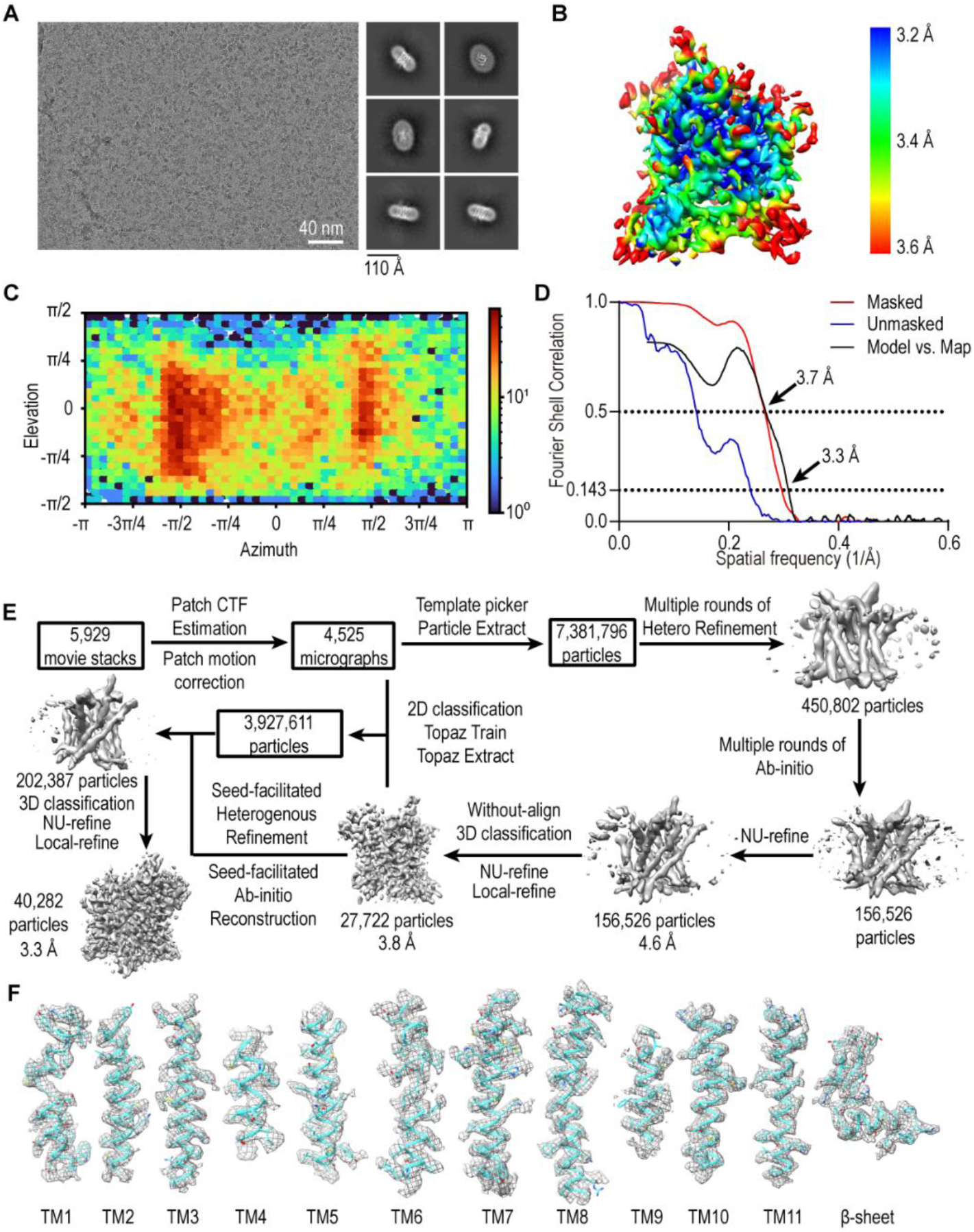
Cryo-EM reconstruction of VIAAT-GABA. **(A)** Representative micrograph (left) (scale bar, 40 nm) and 2D class averages (right). **(B)** Local resolution distribution of VIAAT-GABA. **(C)** Angular distribution of the final reconstruction. **(D)** Gold standard Fourier Shell Correlations (FSC) curves of unmasked map (blue), masked map (red) and model versus map (black), marked with resolutions corresponding to FSC = 0.5 and 0.143. **(E)** Flow chart for cryo-EM data processing of VIAAT-GABA. Further details can be found in the Methods. **(F)** Cryo-EM densities of TM1-11 and the β-sheet in VIAAT-GABA, contoured at 0.3.

**Fig. S5.**
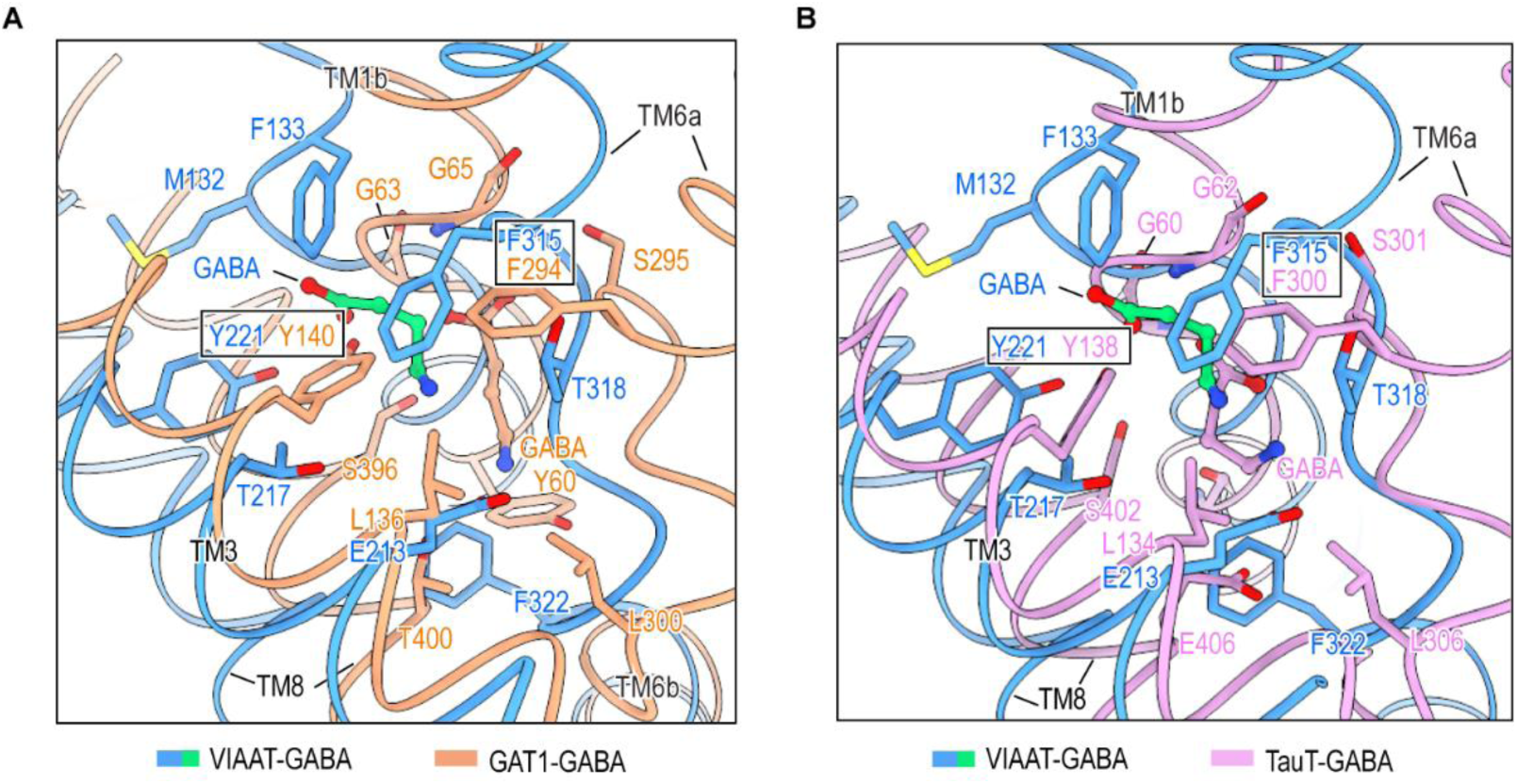
Structural comparison of VIAAT-GABA with GABA-bound GAT1 and TauT. **(A-B)** Superimposition of the binding pocket of VIAAT with those of GAT1 (**A**) and TauT (**B**). GABA-bound VIAAT (blue), GABA-bound GAT1 (light salmon; PDB code 7Y7W), and GABA-bound TauT (plum; PDB code 9JD4) are shown as cartoons. GABA in VIAAT-GABA is shown in green. Two conserved positions corresponding to Tyr221 and Phe315 in VIAAT are boxed.

**Fig. S6.**
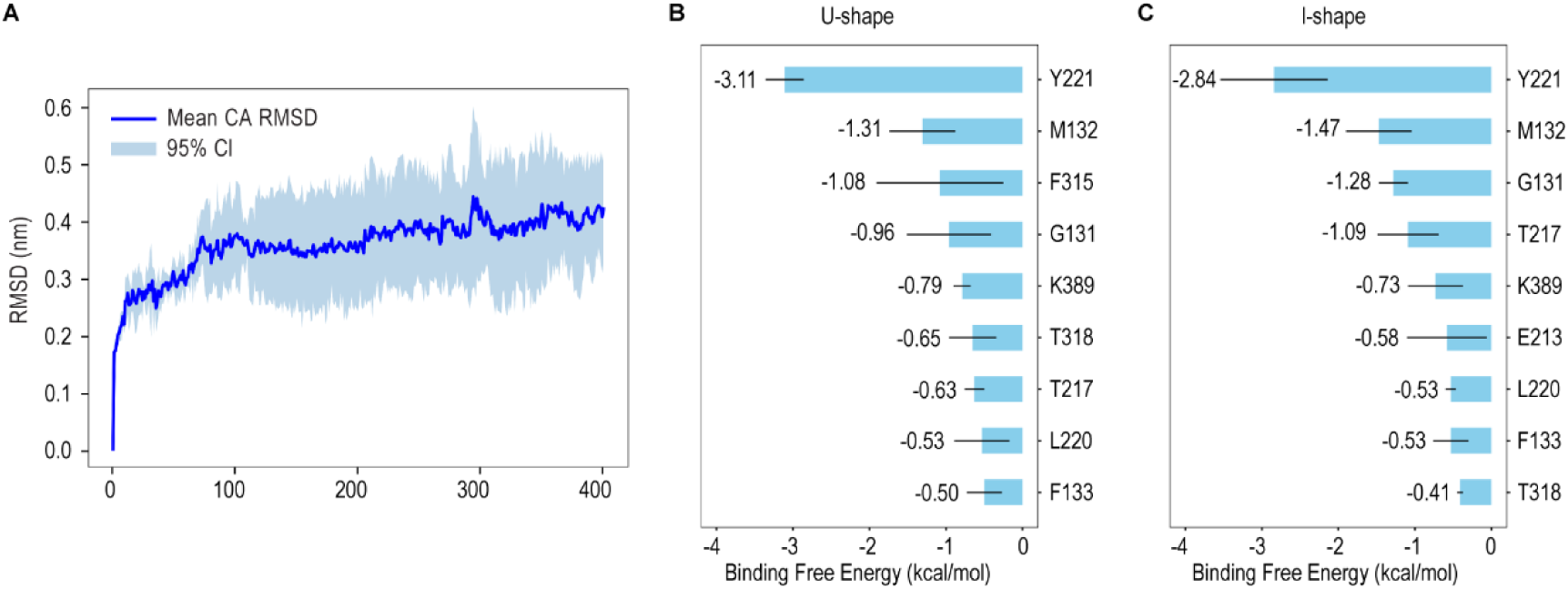
Molecular dynamics simulations of GABA binding to VIAAT. **(A)** Root-mean-square deviation (RMSD) of the VIAAT protein during molecular dynamics simulations. The solid curve represents the average RMSD from three independent replicates, and the shaded region indicates the 95% confidence interval. The protein reached equilibrium after 100 ns in all three replicates. **(B)** Binding free energy contributions of key residues in the U-shape and I-shape binding modes. The top nine residues contributing to binding are shown for each mode, with average values indicated. Horizontal bars represent standard deviations of the binding free energy.

**Fig. S7.**
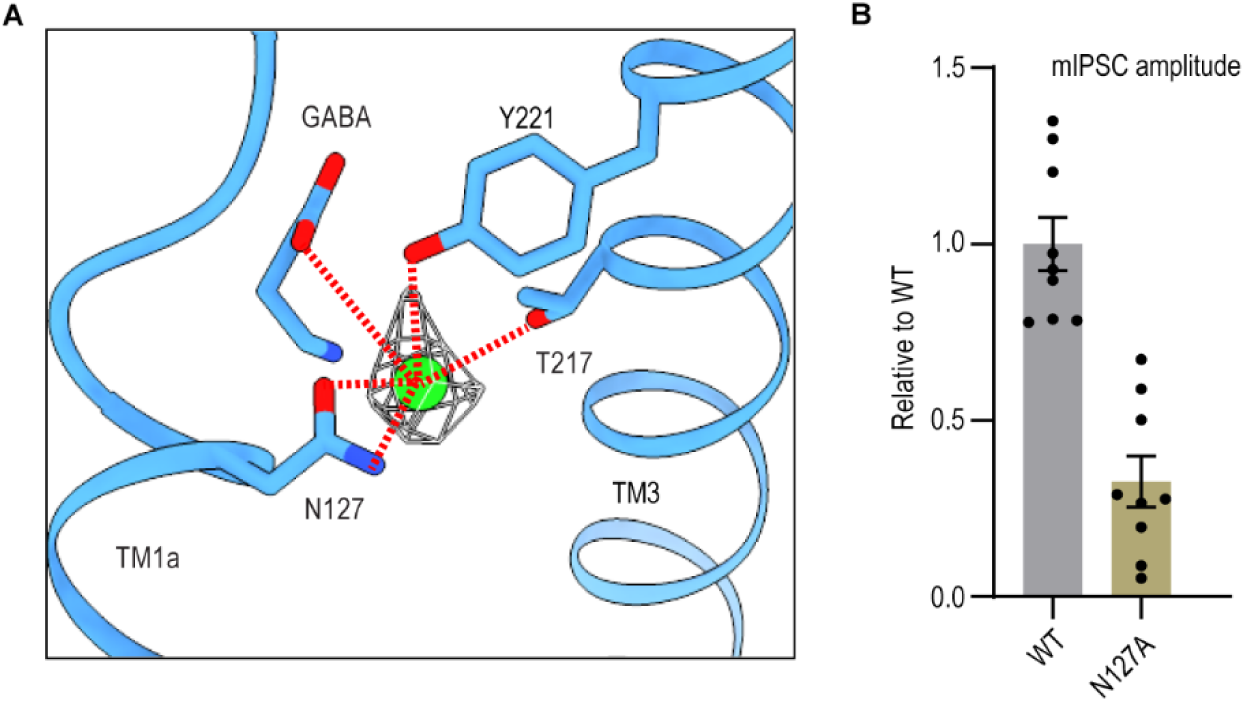
Potential chloride ion binding site. **(A)** Putative coordination of a chloride ion (Cl⁻) in the central binding pocket of VIAAT-GABA. Residues potentially involved in coordination are shown as sticks and labeled. The contour level is 0.35. **(B)** mIPSCs recorded from hippocampal neurons after knockdown of endogenous VIAAT and rescue with WT hVIAAT or its mutants. mIPSC amplitudes were normalized to those of VIAAT-WT (n = 9 cells per condition). Data are shown as mean ± s.e.m.

**Table 1.** Statistics of 3D reconstructions and model refinement.

| **Protein** | VIAAT-apo<br>(PDB-27TQ, EMD-81415) | VIAAT-GABA<br>(PDB-27TR, EMD-81416) |
| --- | --- | --- |
| **Data collection and processing** |  |  |
| Magnification | 81K | 81K |
| Voltage (kV) | 300 | 300 |
| Electron exposure (e<sup>-</sup>/Å<sup>2</sup>) | 50 | 50 |
| Defocus range (μm) | -1.0 ~ -2.0 | -1.0 ~ -2.0 |
| Magnified pixel size (Å) | 1.055 | 1.055 |
| Micrographs | 5,351 | 5,929 |
| Initial particle images (no.) | 4,979,976 | 7,381,796 |
| Final particle images (no.) | 34,411 | 40,282 |
| Symmetry | C1 | C1 |
| Map Resolution (Å) | 3.25 | 3.3 |
| Sharpening B-factor (Å<sup>2</sup>) | 90.6 | 104.3 |
| **Model statistics** |  |  |
| Map CC | 0.65 | 0.67 |
| Number of atoms | 3,271 | 3,177 |
| **R.m.s. deviations** |  |  |
| Bond length (Å) | 0.002 | 0.003 |
| Bond angle (°) | 0.538 | 0.568 |
| **Ramachandran plot** |  |  |
| Favored (%) | 96.69 | 96.62 |
| Allowed (%) | 3.31 | 3.38 |
| Disallowed (%) | 0.00 | 0.00 |
| Rotamers Outlier (%) | 0.00 | 0.00 |
| MolProbity Score | 1.68 | 1.61 |
| Clash score | 8.66 | 7.20 |

